# BeadBuddy: user-friendly, nanometer-scale registration of single-molecule imaging data

**DOI:** 10.1101/2025.11.25.690467

**Authors:** Finnegan T Clark, Peter H Whitney, Néstor Saiz, Angelina Ziarno, Timothée Lionnet

## Abstract

Single-molecule localization microscopy (SMLM) captures nanoscale detail with fluorescence imaging. As large-area cameras and multiplex imaging become standard, chromatic errors are growing more complex, challenging precise analysis. To address this challenge, we present *BeadBuddy*, an open-source, user-friendly software that uses images of fluorescent beads to model and correct 3D, spatially varying chromatic errors. BeadBuddy achieves sub-voxel resolution in DNA Fluorescence In Situ Hybridization (FISH) and is applicable across SMLM modalities.

## Main Text

SMLM has unlocked new insights across diverse fields of study ranging from spatial organization of the genome^1^ to transcription dynamics, neuronal synapsing, and molecular organization of myriad subcellular structures^2^. Multi-channel SMLM requires both accurate localization of diffraction limited spots as well as robust handling of chromatic errors. These errors are introduced by unavoidable aberrations present in every imaging system as well as imperfect alignment of the different imaging channels. Indeed, subcellular distances of interest often span the same length scale as the chromatic errors present in typical microscopes.

Common approaches to chromatic error correction include applying affine transformations constructed from in-situ fiducials or subtracting average inter-channel offsets computed from raw data or fluorescent bead images^34–6^. A key limitation of these approaches is that they treat chromatic errors as invariant across the field of view. In reality, chromatic errors vary as a function of position within the field of view, and these spatial variations become more prominent as the camera sensor size increases^7^. It is therefore critical that error correction methods account for both channel dependence and position dependence within the user’s unique imaging system. Previous work has used an interpolation model to achieve 3D registration of SMLM data to the order of tens of nanometers^8^. Yet, there are still no open-source, flexible and user-friendly tools available to date.

Here, we introduce BeadBuddy, a graphical user interface-based, MATLAB tool for chromatic error correction in multi-channel SMLM datasets. The tool requires minimal user input and no coding experience. We demonstrate its utility in the context of 3D DNA FISH, measuring multi-way genomic interactions at the *PPARG* locus in a human mesenchymal stromal cell adipogenesis model.

Beadbuddy’s workflow begins by imaging a calibration slide of diffraction-limited, commercially available, multifluorescent beads using all color channels relevant to the experiment (Fig. 1a). While each bead fluoresces in all channels, its apparent 3D position varies based on which channel is used to image it. BeadBuddy performs automated 3D subpixel localization of the beads using AIRLOCALIZE to determine bead centroid locations in each channel^9^, then quantifies 3D inter-channel nearest neighbor displacement as a function of x,y position in the field of view. Because of the bead sparsity, the nearest neighbor distance here measures the same bead acquired in different channels, i.e. the chromatic error. BeadBuddy uses a polynomial model to fit the displacement field *F*_*i->ref*_*(x,y) = {dx*_*i->ref*_, *dy*_*i->ref*_, *dz*_*i->ref*_*}* between each non-reference channel and the reference channel (Fig. 1b). The resulting fit functions effectively register all SMLM data to a common reference channel. A graphical user interface walks the user through the process and outputs a table of corrected data as well as useful visualizations.

**Figure 1:**
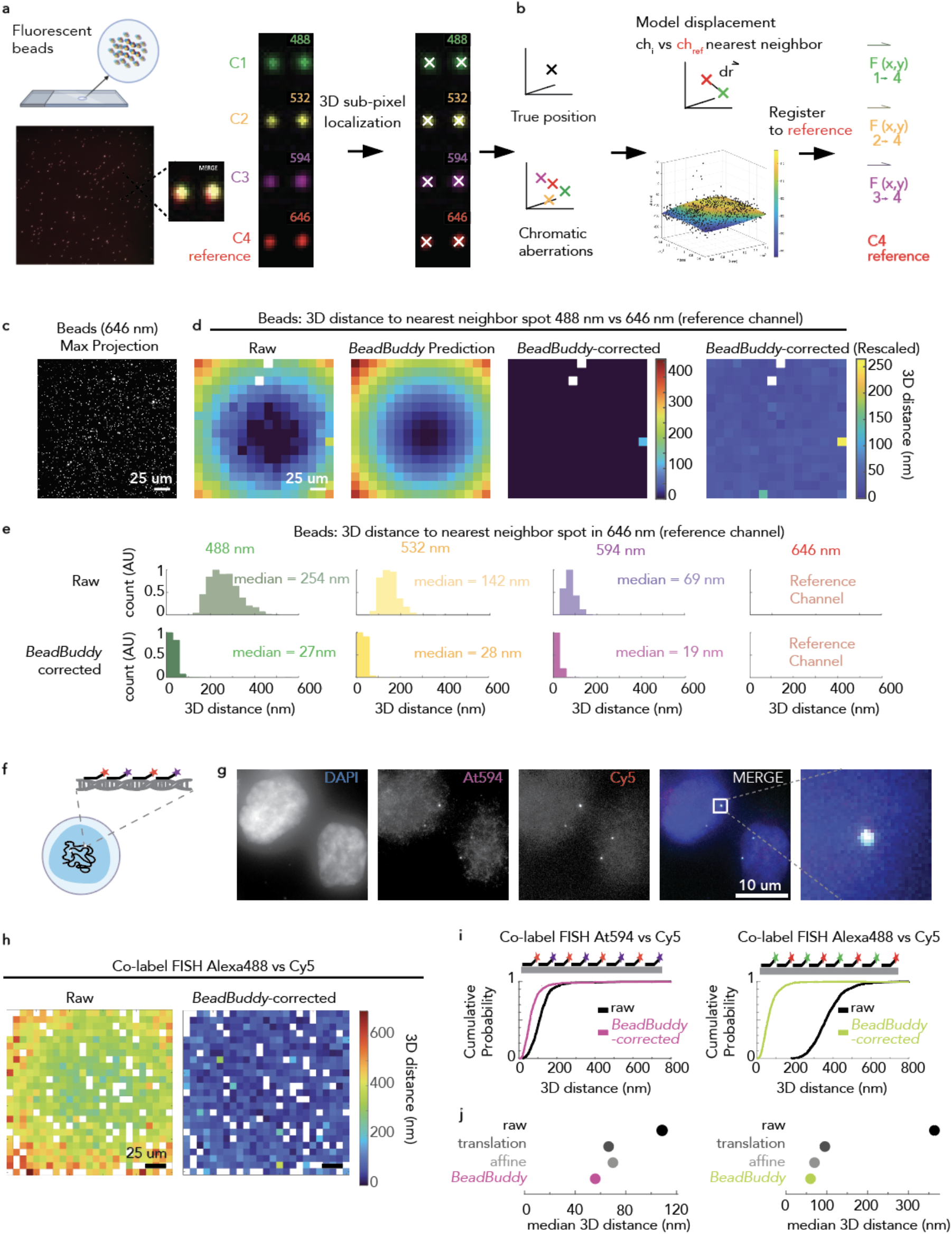
BeadBuddy principle and application to single-molecule imaging data. a) A multi-fluorescent bead calibration slide is imaged in all channels relevant to the experiment. Images are pre-processed and localization of beads is performed using AIRLOCALIZE. b) Nearest neighbor assignment between channels is used to characterize chromatic error. BeadBuddy generates *F*_*i-4*_*(x,y)*, functions that map spot coordinates in each channel *i* to coordinates relative to the chosen reference channel (here C4). c) Example z-max projection (646 nm) from a multi-fluorescent bead slide (distinct from the calibration slide) imaged to quantify chromatic error. d) Heatmaps visualizing the measured 3D distances between each spot in the 488 nm channel and its nearest neighbor in the 646 nm channel (same bead in two channels) as a function of x,y position in the field of view, before and after BeadBuddy. The BeadBuddy prediction panel shows the 3D displacement predicted by BeadBuddy based on a separate calibration. Contrast is enhanced in the rightmost panel for clarity. e) Distributions of measured 3D distances between each channel spot and the nearest 646 nm spot before (raw) and after correction with BeadBuddy. The true physical distance is zero. f) A single genomic locus (*PPARG* downstream enhancer) co-labeled with interspersed Atto594- and Cy5-conjugated oligo DNA FISH probes constitutes a two-color diffraction-limited emitter within a cell. g) Representative image of DNA FISH co-labeled cells as described in (f). h) Genomic locus (*CRISP3*) co-labeled similarly to (f) but with A488 and Cy5. Heatmaps visualizing the 3D distances between each DNA FISH spot in the 488 nm channel and its nearest neighbor in the 646 nm channel as a function of x,y position in the field of view, before and after BeadBuddy. i) Empirical cumulative distribution functions for the inter-channel 3D distances of co-labeled DNA FISH loci before and after correction with BeadBuddy. Atto594 *vs*. Cy5: *PPARG* downstream enhancer. Alexa488 *vs*. Cy5: *CRISP3*. j) BeadBuddy outperforms other common techniques for correction of chromatic error when applied to 3D DNA FISH data from (e-i). Detailed Statistics are presented in Supplementary Table 1.

We first validated BeadBuddy’s performance on fluorescent bead SMLM data, acquiring images in four channels of a wide-field epifluorescence microscope: 488 nm, 532 nm, 594 nm, and 646 nm, with the 646 nm channel arbitrarily chosen as the reference channel (Fig. 1c). The raw chromatic error, quantified as 3D nearest-neighbor distances (reference vs. non-reference channels), varied substantially by channel as well as by position in the field of view (Fig. 1d-e). Once corrected using BeadBuddy, errors were reduced in magnitude and variance (Fig. 1d-e), with corrected 3D median errors in the 19-27 nm range (Fig. 1d,e). This is well below the diffraction limit (usually ∼hundreds of nm) and close to the size of large macromolecular complexes. Additional statistics are listed in Supplementary Table 1.

Next, we validated BeadBuddy in the context of biological samples. To generate two-color, diffraction-limited emitters inside cells, we co-labeled single genomic loci, specifically the *PPARG* downstream enhancer and the *CRISP3* locus, using oligo DNA FISH probes conjugated to two different fluorophores (Fig. 1f,g) ^3,10^. We measured the inter-channel nearest-neighbor distances before and after BeadBuddy correction. BeadBuddy reduced the 3D median error from 109 nm to 59 nm for the Atto594-Cy5 co-label condition and from 363 nm to 58 nm in the Alexa488-Cy5 condition as well as reduced spatial variance in both (Fig 1h,i). We found that BeadBuddy outperformed affine and translational-offset error correction (Fig. 1j).

To confirm that BeadBuddy is effective across diverse imaging platforms, we imaged similar DNA FISH co-labeled samples on a wide-field microscope equipped with three separate cameras and emissions pathways^11^. We observed similar improvements in registration with BeadBuddy (Supplementary Fig.1).

We then set out to measure multi-way chromatin interactions that would otherwise be obscured by chromatic errors. We performed four-channel DNA FISH to characterize the remodeling of chromatin architecture at the *PPARG* locus in a human mesenchymal stromal cell (MSC) model during adipogenesis (Fig. 2a); the expression of the *PPARG2* transcript turns on early during adipogenesis^12,13^.

**Figure 2:**
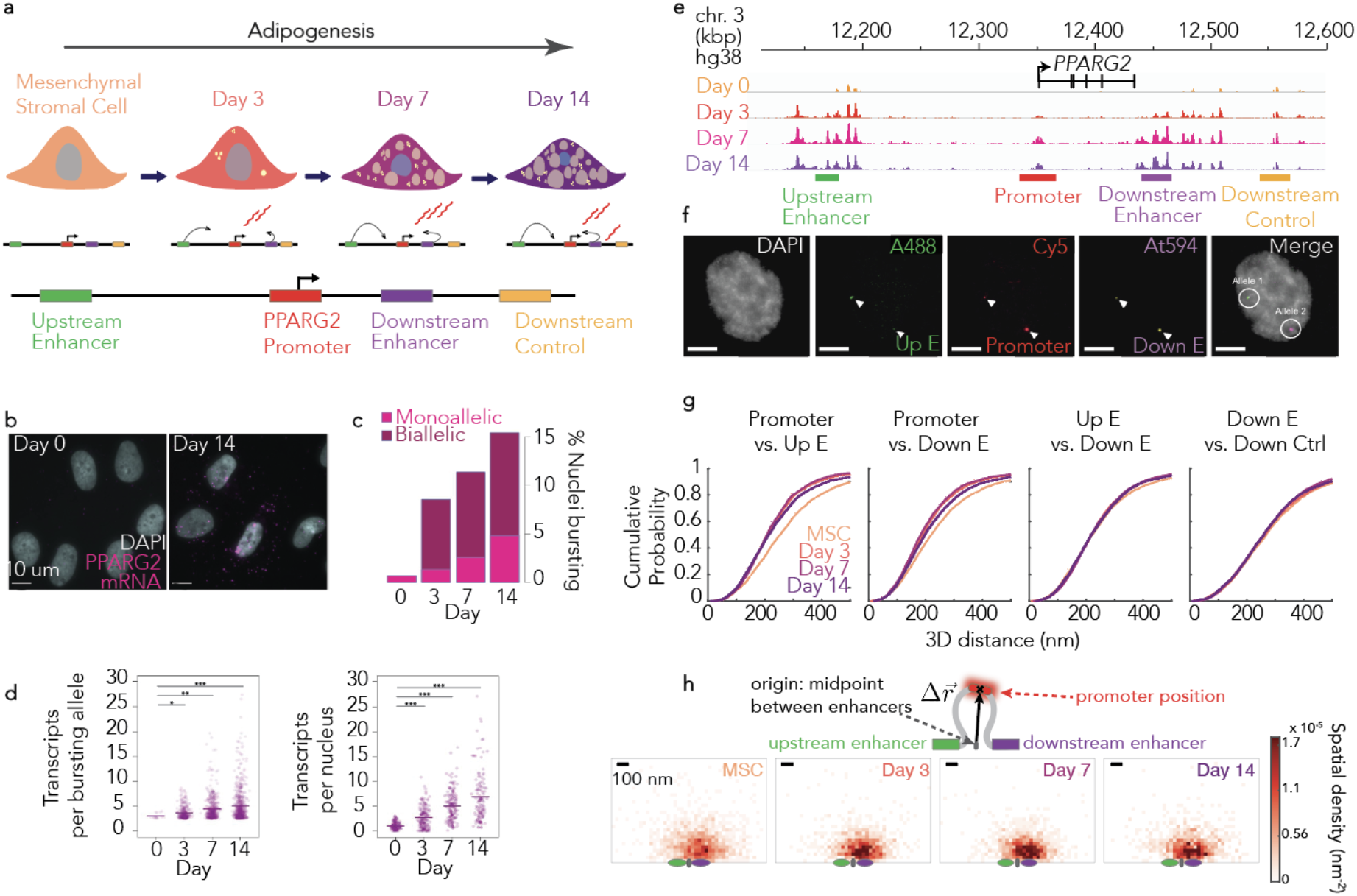
BeadBuddy reveals complex chromatin architecture changes during adipogenesis. a) *In vitro* adipogenesis model. Mesenchymal stromal cells (MSCs) were plated and cultured in either regular or differentiation medium for indicated durations. b) Representative image of *PPARG* RNA FISH in undifferentiated MSCs (Day 0) and adipocytes (Day 14). c) Proportion of nuclei exhibiting bursts of transcription on one (monoallelic) or two of the alleles (biallelic) across four timepoints. d) Left: number of transcript-intensity equivalents per burst at each active transcription site. Right: total number of nuclear transcripts per nucleus. * P < 0.05, ** P <0.01. *** P <0.001. e) Top: H3K27ac ChIP-seq tracks of the *PPARG* locus during adipogenic differentiation (range: 0-30 for all tracks); bottom: Genomic target regions labeled by DNA FISH in four colors. f) Representative maximum intensity projection of DNA FISH data (downstream control locus not shown). g) Pairwise 3D distances between genomic regions of interest measured by DNA FISH and corrected using BeadBuddy. Empirical cumulative distribution functions visualizing changes in distance between specified genomic elements as a function of differentiation timepoint (average of two technical replicates for each timepoint). h) Spatial density heatmaps: each datapoint is an absolute distance of the promoter from the midpoint of the two enhancers on that promoter’s allele. Green and purple ovals represent a distribution of enhancer positions relative to the midpoint (the origin) where each oval’s long axis is 1 standard deviation of enhancer distance from the midpoint. Detailed Statistics are presented in Supplementary Table 1.

We confirmed the transcriptional activation of *PPARG* during differentiation using RNA FISH (Fig 2 b-d). We then simultaneously captured by DNA FISH the relative 3D positions of key genomic elements of the *PPARG* locus - the promoter, two enhancers, and a control region - at various differentiation timepoints (Fig 2e,f). Consistent with increased enhancer-promoter interactions, the distances between each enhancer and the promoter decreased from day 0 to day 7 (upstream enhancer - promoter from 254 ± 11 nm to 208 ± 11 nm; downstream enhancer to promoter from 220 ± 11 to 183 ± 11 nm). Surprisingly, the distance between the two enhancers remained constant (Fig. 2g). To visualize the corresponding 3D architecture, we mapped the distribution of promoter positions relative to the midpoint between the two enhancers (Fig. 2h). This analysis revealed an apparent “reeling in” of the promoter even as the inter-enhancer distances were on average unchanged, consistent with previous findings showing increased confinement of transcribed chromatin regions, including at *PPARG*^*14*^.

BeadBuddy offers a user-friendly and open-source tool to correct complex chromatic errors in multi-channel, SMLM datasets, validated across different imaging systems. BeadBuddy’s input and output files are tables that easily integrate into the user’s pipeline. Application to live imaging and/or other single-molecule localization microscopy modalities require no alterations to the current code. Adoption of BeadBuddy will standardize a critical component of image analysis and improve reproducibility.

## Acknowledgements

P.H.W. is supported by NIH fellowship F32GM153131. T.L. is supported by NIH grants R01AG075272, R01CA260028, R01GM149835 and RM1HG009491.

---

**Supplementary Figure 1.**
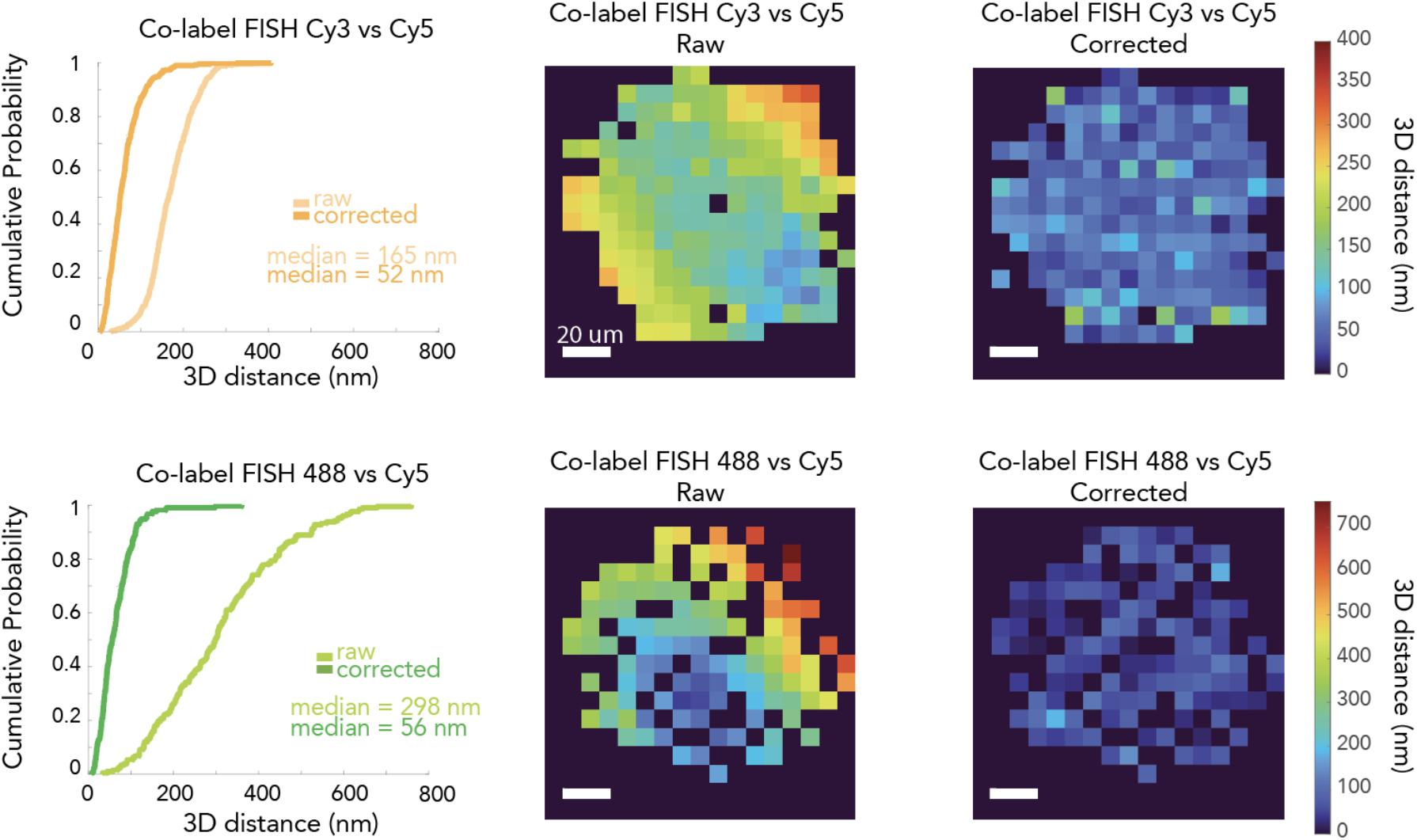
a) Cy3-Cy5 co-labeled *HBB* locus in SEM cells acquired using a multi-camera imaging system with three distinct emission pathways for A488, Cy3, and Cy5 channels. Empirical cumulative distribution function before and after BeadBuddy. Heatmaps visualize the chromatic error profile before and after BeadBuddy. b) Same as (a) but 488-Cy5 co-labeling was used to label the *CRISP3* locus.

**Supplementary Figure 2.**
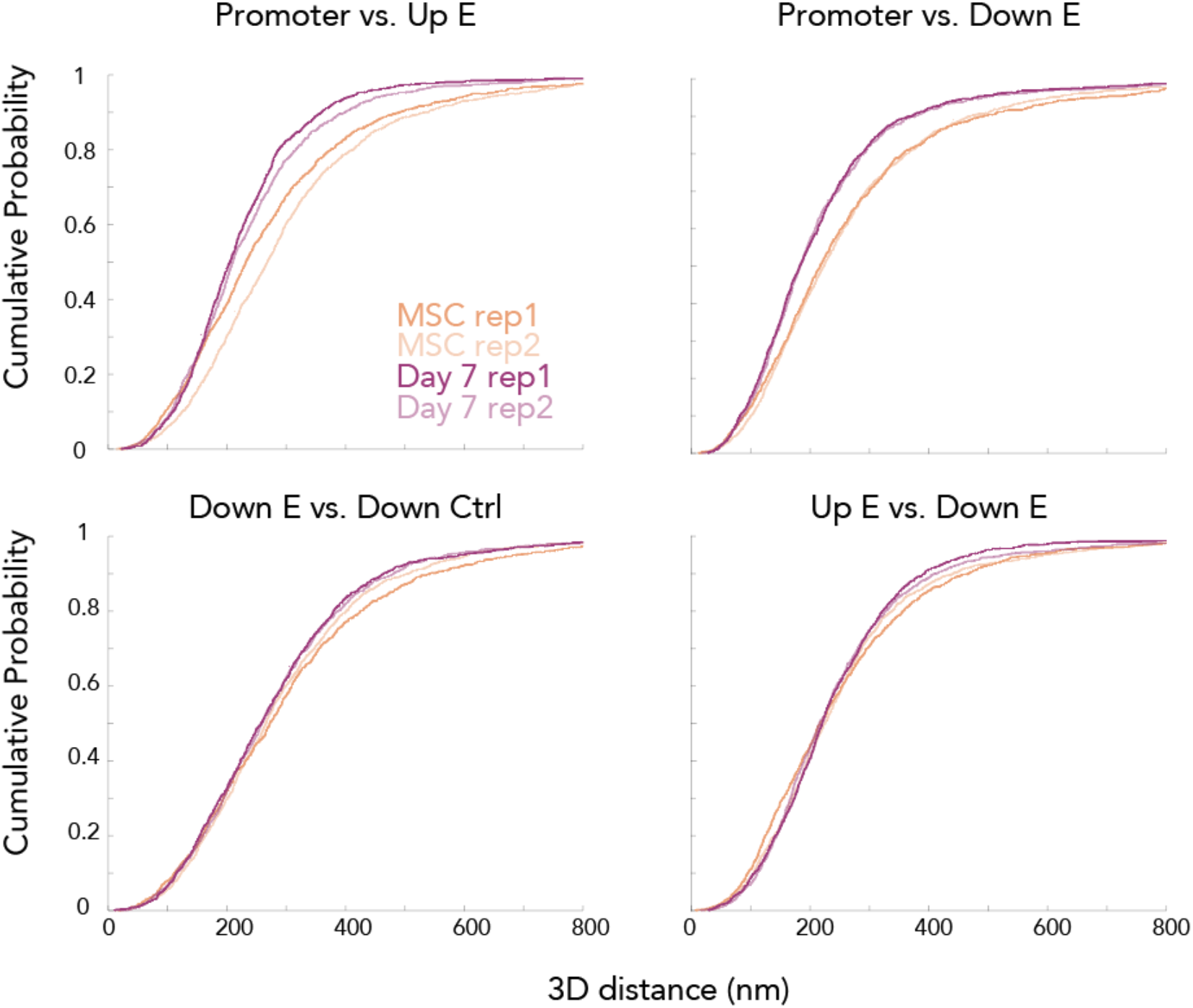
Median 3D distances between genomic targets of interest at the *PPARG* locus during adipogenesis, separated by replicate. Data associated with Fig. (2g). Distances calculated from multi-channel 3D DNA FISH imaging data.

## Materials and Methods

### Cell Culture

#### MSC

Human mesenchymal stromal cells (hMSCs, ATCC: PCS-500-011) were cultured in Dulbecco’s modified Eagle’s medium with Glutamax (DMEM, Thermo: 10-569-044) supplemented with 10% FBS (Avantor Seradigm, VWR 1500-500) and 1X penicillin-streptomycin (pen/strep) (Thermo Scientific 15140122) during expansion and for the undifferentiated condition. Differentiation was performed on 1.5H coverslips (Thorlabs CG15CH) in 6-well plates coated with collagen (Sigma: C3867-1VL). Collagen for coating was diluted 1:50 in 0.02 N acetic acid and coating was performed at 37 C for at least 1 hour. Cells were seeded at approximately 50% confluence (100k cells per well) and allowed 24-48 hrs of normal growth in DMEM. To induce differentiation, media was changed to Minimum Essential Medium α (αMEM, Thermo: 12571063) supplemented with 10% FBS, 1X pen/strep, 250nM Dexamethasone (Sigma: D4902), 500uM IBMX (Cayman Chemical Company: 13347), 1uM Rosiglitazone (Sigma: R2408), and 5mg/ml Insulin (Santa Cruz: sc-360248). Media was changed daily during differentiation and timepoints were collected at 3, 7, and 14 days, with an additional undifferentiated control. Upon collection, cells were washed once with PBS and fixed in fresh 4% PFA in PBS for 10 minutes. After fixation, cells were washed 3 times in PBS and either processed immediately (DNA FISH) or stored at -20 C in 100% ethanol for future processing.

#### HCT116

Human colon carcinoma cells (HCT116, ATCC CCL-247) were cultured in McCoy’s 5A medium (Thermo Scientific 16600082) supplemented with 10% FBS and 1X pen/strep . For DNA FISH, cells were seeded on sterilized 1.5H glass coverslips placed in wells of a 6-well plate (CellTreat), previously coated with a solution of 0.1% gelatin (EMD Millipore ES-006-B) for 10 minutes at room temperature. Cells were seeded at 10^6^ cells/well one day prior to performing DNA FISH and allowed to attach and grow overnight (18-24h).

#### SEM

The SEM acute lymphoblastic leukemia cell line was a gift from the Caroll lab at NYU Langone. Cells were cultured in RPMI media (ThermoFisher 11875093) supplemented with 10% FBS and 1X pen/strep in T-75 flasks (Corning 431463). Cells were maintained at 10^6 / mL and were split approximately 1:5 every 3 days. Cells were concentrated to 10^7 / mL in PBS. A 30 uL droplet of cells was allowed to sediment 5 minutes and adhere to 1.5H coverslips coated with 0.01% poly-L-lysine . The coverslip was gently washed once in PBS and then fixed with 4% PFA in PBS for 10 minutes. After fixation, cells were washed 3 times in PBS and processed immediately (DNA FISH).

### Preparation of Bead Calibration slide

A precision #1.5H coverslip was coated with 0.1% Poly-L-Lysine (Sigma P8920) by incubating it at room temperature for 5 minutes, rinsing three times with DI H2O, and allowing it to dry at room temperature. A suspension of fluorescent beads (TetraSpeck 0.1 um microspheres, ThermoFisher T7279) was diluted 1/100 in DI H2O after thoroughly mixing by vortexing and pipetting in order to reduce clumping of beads. 20 uL of the diluted bead suspension was spread over the coated coverslip and allowed to dry out at 37 C. The coverslip was then mounted onto a microscopy slide using a drop of ProLong Gold mounting medium (ThermoFisher P36930) and allowed to cure in the dark for 48 hours. The coverslip was then sealed with nail polish and stored at RT indefinitely.

### DNA FISH Probe design and synthesis

#### Protocols based on ^3,15,16^ with some modifications

Oligo probes targeting the *HBB* locus and the *CRISP3* locus were ordered from Daicel Arbor Biosciences using the myTag Custom Probe product. Oligo probes targeting the *PPARG* locus were designed using paintSHOP ^10^ and were then ordered from IDT using the oPools Oligo Pools product. Non-commercial DNA FISH Probe sequences are listed in Supplementary Table 2.

#### Limited cycle PCR amplification of probe library

Single-stranded DNA (ssDNA) probe libraries (custom oPools or myTags) were diluted to ∼0.1 ng/μL and amplified using KAPA HiFi HotStart ReadyMix (Roche, Cat. No. 50-196-5217) with 100 μM forward and reverse primers specific to each probeset. PCR was performed with the following conditions: 95 °C for 3 min; 20 cycles of 98 °C for 20 s, 56 °C for 15 s, and 72 °C for 30 s; followed by a final 72 °C extension for 30 s. Then a debubbling mix (10 uL KapaHiFi Hotstart Readymix, 1.2 uL PCR primer mix, and 8.8 uL nuclease-free water) was added and the same PCR program was repeated 2x. PCR products (∼100 bp) were visualized on a 1.8% agarose gel and purified using the QIAquick PCR Purification Kit (Qiagen, Cat. No. 28104).

#### In vitro transcription (IVT)

Purified double-stranded DNA templates were transcribed using the HiScribe T7 High Yield RNA Synthesis Kit (NEB, Cat. No. E2040S) at 37 °C for 4–16 h (saturating product). RNA was purified using the NucleoSpin RNA Clean-up Kit (Macherey-Nagel, Cat. No. 740948.50) and quantified on a NanoDrop spectrophotometer.

#### Reverse transcription and probe labeling

Complementary DNA (cDNA) was synthesized using SuperScript IV Reverse Transcriptase (Thermo Fisher Scientific, Cat. No. 18090010) with 52 μg RNA template, 1 mM directly conjugated, fluorescently labeled RT primer, and 10 mM dNTPs (NEB, Cat. No. N0447S). Reactions were incubated at 50 °C for 4 h. Unincorporated primers were digested with Exonuclease I (NEB, Cat. No. M0293S) at 37 °C for 15 min, followed by heat inactivation in 0.5 M EDTA, pH 8.0, at 80 °C for 20 min to generate RNA:DNA hybrids.

#### Purification and RNA digestion

RNA:DNA hybrids were purified using the Zymo Quick-RNA Miniprep Kit (Zymo Research, Cat. No. R1054). Residual RNA was digested using RNase H (NEB, Cat. No. M0297S) and RNase A (Thermo Scientific, Cat. No. EN0531) using the program (37 °C × 2 h; 70 °C × 20 min; 50 °C × 60 min). The resulting single-stranded DNA (ssDNA) probes were purified using the Zymo Quick-RNA Miniprep Kit and eluted in 100 μL nuclease-free water.

#### Quantification and labeling efficiency

Labeled ssDNA probe concentrations were determined using the NanoDrop Microarray setting (Thermo Fisher). Concentrations were converted from ng/μL to pmol/μL using a molecular weight of 0.047 pmol/ng for 66 bp probes. Labeling efficiency was calculated as the ratio of fluorophore to ssDNA concentration, with typical efficiencies exceeding 90%.

### Oligo DNA FISH

Protocols based on ^3,15,16^ with some modifications. All steps were performed at room temperature (RT) unless otherwise indicated.

#### Sample preparation

Cells grown on #1.5H coverslips were fixed in 4% paraformaldehyde in PBS, rinsed three times in PBS, and processed on the day of fixation in 6-well plates. Cells were permeabilized in 0.5% Triton X-100/PBS (vol/vol) for 10 min, rinsed twice for 3 min in PBS, then incubated in 0.1 N HCl for 5 min (time-sensitive step). Following three 1-min rinses in 2× SSC and three 3-min rinses in PBS, samples were incubated with RNase A 10 ug/mL in PBS (vol/vol) (ThermoFisher EN0531) at 37 °C for 1 h in a humid chamber.

#### Pre-hybridization and probe mix preparation

Samples were pre-equilibrated in pre-warmed Hybridization buffer #1 (0.1% Tween-20 (vol/vol), 50% formamide (vol/vol) and 2xSSC (final conc) in UltraPure H2O (ThermoFisher 10977015)) at 37 °C for 30 min. Meanwhile, to prepare probe mix, FISH probes were concentrated and resuspended in Hybridization buffer #2 (50% formamide (vol/vol), 0.1% Tween-20 (vol/vol), 10% (vol/vol) dextran sulfate and 2× SSC (final conc) in UltraPure H2O) at a final volume of 20 µL per coverslip, with probe volume comprising <5% of the total mix. Each 20 µL of probe mix contained 4 pmol of each probe set. Probe mix was denatured for 5 min at 85 °C in the dark, then cooled on ice for 5 min.

#### Denaturation

Approximately 20 µL of the probe mix was pipetted onto a microscope slide (avoid bubbles), and the sample was inverted onto the drop and sealed with rubber cement. After drying in the dark for 5 min, samples were denatured for 5 min at 85 °C on an aluminum heating block inverted and submerged (except for the top 0.5 cm) in a water bath (critical step; confirm temperature and maintain humidity). Slides were transferred to a humid and light-tight tray, then hybridized overnight (>16 h).

#### Post-hybridization washes

Coverslips were floated off slides using a reservoir of 2× SSC (rubber cement should easily peel away). Coverslips were transferred to 6-well plates containing probe wash buffer and incubated at 42 °C for 50 min in a humid chamber; this step was repeated once with fresh buffer. Samples were rinsed twice for 5 min in 5× SSCT, followed by a 5-min PBS rinse. Nuclei were counterstained with Hoechst for 5 min, then washed in PBS until mounting (make sure cells do not dry out).

### RNA FISH

RNA FISH probes targeting the *PPARG2* transcript were designed by and ordered from Molecular Instruments.

Following fixation in freshly prepared 4% paraformaldehyde in PBS, cells on coverslips were washed 3x in PBS then permeabilized in 0.2% Triton X-100 in PBS for 10 minutes. Coverslips were then transferred to Molecular Instruments Hybridization Buffer at 37 C and pre-hybridized for 30 minutes. Probe hybridization was then performed overnight at 37 C with 1 pmol of PPARG primary probes in Molecular Instruments Hybridization Buffer. After washing in 3x Molecular Instruments Probe Wash Buffer at 37 C, one final wash of 5xSSCT was done at room temperature before pre-amplifying in Molecular Instruments Amplification Buffer for 30 minutes at room temperature. Probe amplification was carried out with denatured B2 amplifier secondary probes (stock concentration, 3µM) diluted 1:50 in Amplification Buffer. Amplification time was kept to 1 hour to ensure linear amplification to enable single molecule quantification. Coverslips were then washed 4x in 5xSSCT followed by 2x wash in PBST (0.1% Tween 20 in PBS).

Finally, cells were stained with Hoechst diluted 1:3000 in PBS, and washed with PBS once before mounting in ProLong Gold mounting media.

### Image acquisition

Data was acquired using two custom-built microscopes. The single-camera epifluorescence microscope consists of an epi-fluorescent Nikon Ti-E microscope controlled by Micro-Manager ^17,18^. Samples were imaged using five channels excited by lasers emitting respectively at 405 nm (DAPI), 488 nm (Alexa Fluor 488), 532 nm (Cy3), 594 nm (Atto Fluor 594) and 637 nm (Cy5), and imaged through a 60x oil-immersion Olympus UPlanXApo lens (NA=1.4) onto a Photometrics Prime BSI Express camera using a custom tube lens. Image stacks were captured per sample with an effective voxel size 98 x 98 x 250 nm.

The three-camera epifluorescence microscope is built around a RAMM frame (Applied Scientific Instrumentation) controlled by Micro-Manager, following a prior design ^11^. Samples were imaged using four channels excited by lasers emitting respectively at 405 nm (DAPI), 488 nm (Alexa Fluor 488), 561 nm (Cy3), and 637 nm (Cy5), and imaged through a Nikon CFI SR HP Plan Apo 100x oil-immersion lens (NA=1.35) onto three Photometrics Prime 95B cameras. Image stacks were captured with an effective voxel size 109 x 109 x 250 nm.

Calibration bead images were acquired as follows: before imaging, the microscope was turned on and allowed to thermalize for 2.5 h to avoid position drift induced by thermal expansion of the system components. To thoroughly sample the x,y focal plane, we acquired an x,y grid of Z-stacks in all relevant channels, aiming for a 3x3 grid of Z-stacks with ∼97% overlap between adjacent tiles in the grid and using 250 µm Z-steps (typically 200 - 1000 beads per field of view).

Following acquisition of the bead images, DNA FISH slides were imaged in the same session as a tile-scan using the Micro-Manager MDA and JAF(TB) autofocus tool ensuring the entire nucleus was contained within the stack using 250 µm z-steps.

RNA FISH Coverslips were imaged on the single-camera epifluorescence microscope. We captured approximately 120 fields of view per condition over two technical replicates.

### BeadBuddy Analysis

Calibration bead images were first pre-processed using MIJI to call FIJI from MATLAB as follows: hyperstacks were split by channel, and the {*x*_*i*_, *y*_*i*_, *z*_*i*_*}* coordinates of the center of the image of each bead were localized in each channel *C*_*i*_ separately using AIRLOCALIZE^9^.

Localization intensity thresholds were determined by coarsely matching AIRLOCALIZE output counts to an approximate count of binarized z-max projections of beads.

Sets of coordinates belonging to images of the same bead in different channels were matched using a mutual nearest neighbor criterion: we first computed the all-by-all 3D euclidean distance matrix *dr =* || ***r***_***i***_ *-* ***r***_***ref***_ || between each bead image in the non-reference channel *C*_*i*_ and each bead image in the reference channel. Then, the minimum distance across each row and each column was computed and reciprocal nearest neighbors were paired together to filter out beads that were not detected in all channels.

For each pair of nearest neighbor coordinate sets, chromatic errors *dx*_*i->ref*_ *= x*_*i*_ *-x*_*ref*_, *dy*_*i->ref*_ *= y*_*i*_ *-y*_*ref*_, *dz*_*i->ref*_ *= z*_*i*_ *-z*_*ref*_ were computed. MATLAB’s Curve Fitting Toolbox ***fit*** function with **fitType = ‘poly33’ and ‘Normalize’ = ‘on’** was used to generate a third-degree polynomial surface of best fit, **F**_**i->Ref**_**(X**,**Y)**.

The fit function **F**_**i->Ref**_**(X**,**Y)** was eventually applied to the user’s DNA FISH (or any other type of localization data) from the corresponding non-reference channel *i* (2D or 3D coordinates), to register the measured positions from that channel onto the reference channel and thus minimize chromatic errors.

Additional statistics are listed in Supplementary Table 1.

### DNA FISH Analysis

Images were pre-processed using a custom MATLAB pipeline that called FIJI through MIJI to split hyperstacks and organize files by channel and field of view. Nuclear segmentation was performed on z-maximum projections of the nuclear stain using Cellpose 2.0^19^ with the pre-trained “nuclei” model. Optimal nuclear diameter parameter varied per condition. DNA FISH spots were localized with AIRLOCALIZE^9^ and intersected with the corresponding nuclear masks. A three-dimensional mutual nearest-neighbor analysis was then used to associate DNA FISH spots belonging to the same allele. Quality-control criteria were applied such that cells were excluded if they contained fewer than one or more than four spots per channel. For the four-color *PPARG* DNA FISH experiments, alleles were retained only if at least three of the four genomic elements were successfully detected.

### RNA FISH Analysis

Nuclear segmentation was performed using Cellpose 2.0^19^ using the pre-trained “nuclei” model with nuclei diameter set at 150 px. Transcripts were called using the MATLAB program AIRLOCALIZE^9^. A custom R script was used to assign transcripts to individual nuclei by intersecting the xy coordinates of spot calls with nuclear masks. The number of nascent transcripts was calculated by computing the median fluorescence intensity of all transcript spots, normalizing all spot intensities by this value. Spots with a normalized intensity value >2.5 transcripts were labeled as nascent spots. Monoallelic or biallelic bursting cells were determined by a nucleus containing either one or two nascent spots respectively. Statistical testing for transcripts per burst and transcripts per nucleus performed with two-sided Wilcoxon ranksum test.

## Code availability

Matlab code and detailed instructions for Beadbuddy are deposited at: https://github.com/timotheelionnet/BeadBuddy. Demonstration dataset available at https://zenodo.org/records/17605122.

